# Shared and distinct brain regions targeted for immediate early gene expression by ketamine and psilocybin

**DOI:** 10.1101/2022.03.18.484437

**Authors:** Pasha A. Davoudian, Ling-Xiao Shao, Alex C. Kwan

## Abstract

Psilocybin is a psychedelic with therapeutic potential. While there is growing evidence that psilocybin exerts its beneficial effects through enhancing neural plasticity, the exact brain regions involved are not completely understood. Determining the impact of psilocybin on plasticity-related gene expression throughout the brain can broaden our understanding of the neural circuits involved in psychedelic-evoked neural plasticity. In this study, whole-brain serial two-photon microscopy and light sheet microscopy were employed to map the expression of the immediate early gene, c-Fos, in male and female mice. The drug-induced c-Fos expression following psilocybin administration was compared to that of subanesthetic ketamine and saline control. Psilocybin and ketamine produced acutely comparable elevations in c-Fos expression in numerous brain regions, including anterior cingulate cortex, locus coeruleus, primary visual cortex, central and basolateral amygdala, medial and lateral habenula, and claustrum. Select regions exhibited drug-preferential differences, such as dorsal raphe and insular cortex for psilocybin and the CA1 subfield of hippocampus for ketamine. To gain insights into the contributions of receptors and cell types, the c-Fos expression maps were related to brain-wide *in situ* hybridization data. The transcript analyses showed that the endogenous levels of *Grin2a* and *Grin2b* are predictive of whether a cortical region is sensitive to drug-evoked neural plasticity for both ketamine and psilocybin. Collectively, the systematic mapping approach produced an unbiased list of brain regions impacted by psilocybin and ketamine. The data are a resource that highlights previously underappreciated regions for future investigations. Furthermore, the robust relationships between drug-evoked c-Fos expression and endogenous transcript distributions suggest glutamatergic receptors as a potential convergent target for how psilocybin and ketamine produce their rapid-acting and long-lasting therapeutic effects.

## INTRODUCTION

Psychedelic compounds produce profound changes in states of perception and cognition ^1,2^. These compounds have been studied for their potential therapeutic effect for a variety of psychiatric conditions ^3^. In particular, psilocybin has reemerged recently with several promising early-phase clinical trials for the rapid and sustained treatment of depression ^4–8^. These results have led to an explosion of clinical trials to test the efficacy of psilocybin and other psychedelics as treatment for mental illnesses.

The therapeutic benefits of psychedelics are presumed to depend on neural plasticity ^9–11^. Most recent research to study psychedelics-induced neural plasticity has focused on the neocortex and hippocampus ^12–19^. However, as the compound is delivered systemically, many other regions in the brain can also potentially be responsive to psychedelic administration. Indeed, early work in rodents has shown strong responses to psychedelics in several subcortical nuclei. For example, the dorsal raphe, a key source of serotonin for the forebrain, exhibited a cessation of spiking activity following the administration of lysergic acid diethylamide (LSD) and other psychedelic compounds ^20,21^. Other studies demonstrated increased neural activity of the locus coeruleus in response to peripheral stimuli following ergoline and phenethylamine administration ^22,23^. Therefore, there is incentive to explore the entire brain to illuminate the neural circuits mediating the actions of psychedelics.

Immediate early genes such as c-Fos provide a window into the plasticity mechanisms evoked by a variety of stimuli ^24–26^. Transcription is activated in neurons rapidly within minutes of stimulation, which could be due to spiking activity, but is known to also arise from exposure to growth factors ^27^ and can be pharmacologically induced ^28,29^. Importantly, immediate early genes are thought to mediate key steps in protein synthesis, synaptic potentiation, and structural plasticity ^30,31^. Not surprisingly, given the role of immediate early genes in neural plasticity, psychedelics have been demonstrated to increase expression of c-Fos when measured as transcripts in specific regions such as the frontal cortex, hippocampus, and midbrain ^32,33^, and protein in the anterior cingulate cortex, paraventricular nucleus, bed nucleus of the stria terminalis, and central amygdala ^34–38^. However, these studies focused on pre-determined brain regions for analysis. New technologies such as serial two-photon microscopy and light sheet microscopy enabled whole-brain mapping of c-Fos expression ^39–42^. In this study, we leveraged these technologies to map the brain-wide distribution of c-Fos protein expression following the administration of psilocybin, comparing to the fast-acting antidepressant ketamine and saline controls.

## RESULTS

### Whole-brain imaging of c-Fos expression

Mice received either saline (10 mL/kg, i.p.; n = 4 males, 4 females), ketamine (10 mg/kg, i.p.; n = 4 males, 4 females), or psilocybin (1 mg/kg, i.p.; n = 4 males, 4 females).The psilocybin dose of 1 mg/kg was chosen because of prior work demonstrating that this dose is sufficient to induce robust head-twitch response as well as enduring neural plasticity in the medial frontal cortex and hippocampus ^12,13^. The ketamine dose of 10 mg/kg was chosen because of prior work demonstrating that this subanesthetic dose is sufficient to induce spinogenesis in the frontal cortex and alleviate stress-induced behavioral deficits ^43,44^. To measure the impact of these drugs on the whole-brain expression of the plasticity-related immediate early gene c-Fos, we used two complementary imaging methods. First, we used the *cfos^GFP^* transgenic mouse ^45,46^ with the brain harvested 3.5 hours after drug administration (**Figure 1A**), because prior work demonstrated that the fluorescent signal, arising from a two-hour half-life GFP associated with the c-Fos promoter, peaked at this timepoint ^39^. The fixed brain was imaged with serial two-photon microscopy (**Figure 1B**). This approach allowed us to examine GFP expression in the whole brain with micron-scale resolution (**Figure 1C**). The strength of this method is that the tissue does not need to be cleared, and therefore minimizes distortion. Second, we used the C57BL/6J mouse with the brain collected 2 hours after drug administration (**Figure 1D**), because endogenous c-Fos protein expression peaks at this time ^47,48^. Whole-brain clearing and immunohistochemistry were used to label the endogenous c-Fos protein. The cleared brain was imaged using light sheet microscopy (**Figure 1E**). This latter approach allowed for visualization of c-Fos protein expression at a similar micron-scale resolution (**Figure 1F**). The strength of this method is that imaging more rapid, thus axial sampling can be superior and the whole brain is sampled. Moreover, antibodies tag endogenously produced c-Fos proteins, which avoids potential confounds in using mutant animals where the transgene expression may not reflect endogenous c-Fos levels ^49,50^. Knowing the strengths of each method, the use of both techniques allowed us to generate complementary datasets to examine drug-evoked c-Fos expression levels.

**Figure 1.**
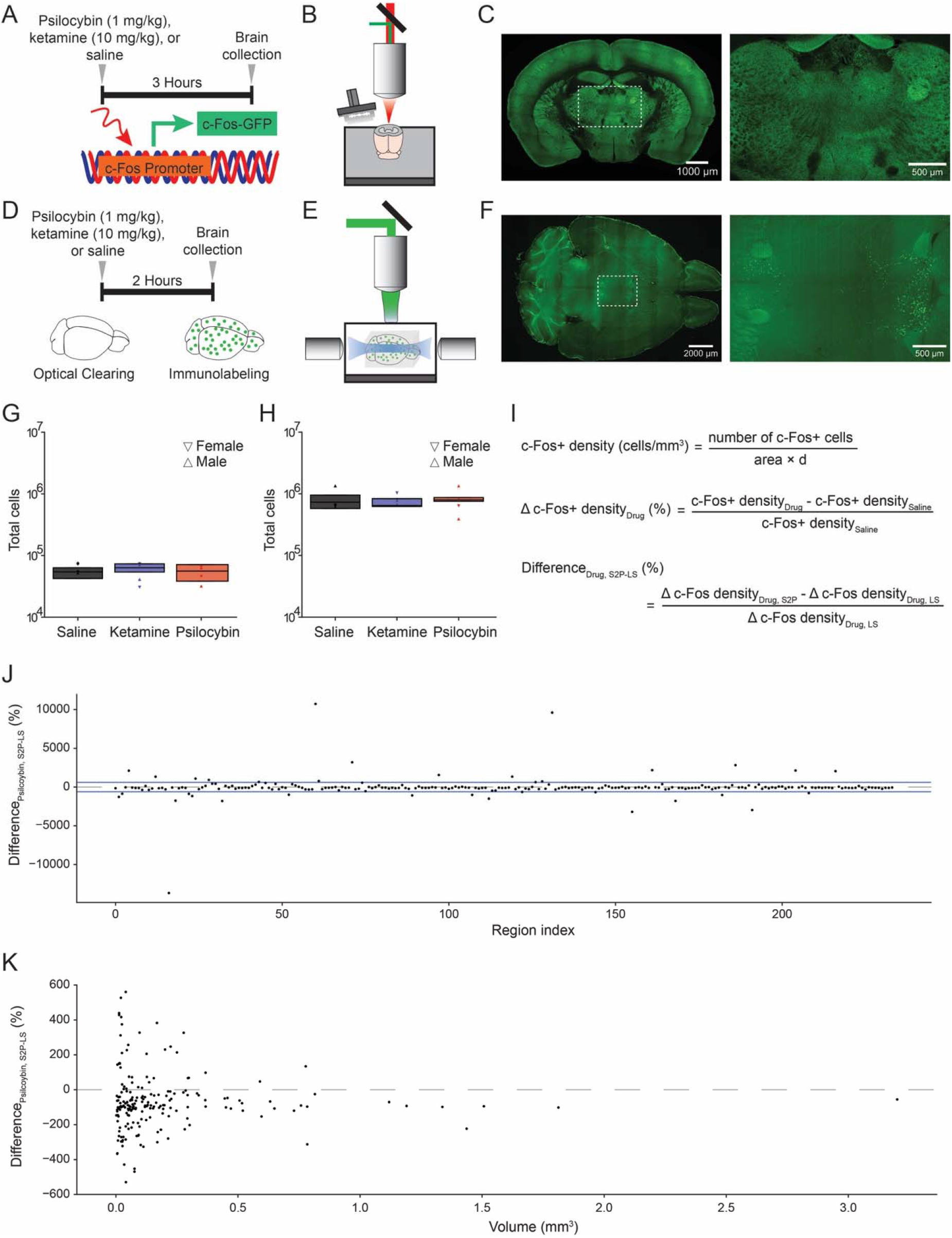
Whole-brain mapping of drug-induced c-Fos expression. **(A)** Transgenic c-Fos-GFP mice were injected with either saline, ketamine (10 mg/kg), or psilocybin (1 mg/kg) at 3 hours before sacrifice and collection of brains (*n* = 4 per condition). **(B)** Schematic of the serial two-photon microscope setup. **(C)** Left: tiled image of a coronal block-face of a brain from c-Fos-GFP mouse. Right: zoomed in view demonstrating expression of c-Fos puncta in neurons. **(D)** C57/BL6 mice were injected with either saline, ketamine (10 mg/kg), or psilocybin (1 mg/kg) at 2 hours before sacrifice and collection of brains (*n* = 4 per condition). Brains were cleared and then immunolabeled with antibody against c-Fos protein. **(E)** Schematic of the light sheet microscope setup. **(F)** Left: image of a horizontal plane of a cleared mouse brain labeled with c-Fos antibody. Right: zoomed in view demonstrating expression of c-Fos puncta in neurons of the cortex. **(G)** Total number of c-Fos+ cells detected in the entire brain across different drug conditions using serial two-photon microscopy. Symbol, individual animal. Box plot shows the median, 25^th^ and 75^th^ percentiles. **(H)** Total number of c-Fos+ cells detected in the entire brain across different drug conditions using light sheet microscopy. Symbol, individual animal. Box plot shows the median, 25^th^ and 75^th^ percentiles. **(I)** Top: formula to calculate c-Fos+ cell density for a region. Middle: formula to calculate the change in c-Fos+ cell density due to drug compared to saline. Bottom: formula to calculate difference in drug-evoked change in c-Fos+ cell density between the two imaging modalities. S2P, serial two-photon microscopy. LS, light sheet microscopy. (**J**) Difference in psilocybin-evoked change in c-Fos+ cell density between the two imaging modalities, plotted by brain region. Region indices are listed in Supplementary Table 1. Blue lines, threshold for exclusion. Dashed line, zero percent difference. (**K**) Difference in psilocybin-evoked change in c-Fos+ cell density between the two imaging modalities, plotted as a function of the volume of brain region as assessed by serial two-photon microscopy. Dashed line, zero percent difference.

### Mapping drug-induced differences in c-Fos expression

To compare between the serial two-photon and light sheet imaging approaches, we first examined the number of c-Fos+ cells in the brain. For both approaches, c-Fos+ cells were identified using automated procedures based on machine learning (see Methods and Materials). We did not detect any difference in the total number of c-Fos+ cells across vehicle and drug treatment conditions (serial two-photon: *P* = 0.6, **Figure 1G**; light sheet: *P* = 1, **Figure 1H**; one-way ANOVA). The serial two-photon approach yielded significantly fewer c-Fos+ cell count than light sheet imaging (*P* = 9 × 10^−8^, two-sided *t*-test; n = 12, 12), which was expected because we used a coarser axial sampling step size in serial two-photon microscopy (100 μm) than light sheet microscopy (4 μm). We split the data by sex and did not detect difference in total c-Fos+ cell count between males and females (saline: *P* = 0.9, ketamine: *P* = 0.7, psilocybin: *P* = 0. 8, two-sided *t*-tests; n = 4, 4 each), however the current study was not powered to detect a sex difference.

Next, we analyzed the density of c-Fos+ cells in each brain region (**Figure 1I**). We followed the guidance of Allen Mouse Brain Common Coordinate Framework, which identified 316 “summary structures” as the basis set for rodent brain parcellation ^51^. First, we assessed potential differences in results generated by the two imaging methods. We started with 234 regions that had a minimum of 10 c-Fos+ cells in each brain across all drug and imaging conditions, because some regions were small and not sampled adequately by serial two-photon microscopy. We then calculated the percent difference change in c-Fos+ cell density between drug and saline (**Figure 1I**). For psilocybin, we note that although drug-evoked changes in c-Fos+ cell density were mostly comparable across the two imaging approaches, there were a subset of regions that showed large differences (**Figure 1J, Supplementary Table 1**). Plotting based on each region’s volume revealed that the smaller regions tend to show larger discordance between two methods, and serial two-photon imaging often under-reports the drug-evoked changes seen in light sheet imaging (**Figure 1K**). The disagreements between the two imaging methods likely arise from differences in sampling density, which has been computationally modelled by another study showing that methods with lower sampling rate yield more variable results ^52^. Another reason for differences is that serial two-photon microscopy produced coronal images (**Figure 1B, C**) whereas light sheet microscopy generated transverse images (**Figure 1E, F**), therefore each can have more favorable sampling for certain brain regions depending on the region’s orientation in the brain. For these reasons, we took away 35 regions that had substantial variations across the two methods (threshold indicated by blue lines in **Fig. 1J**; see Methods and Materials). This yielded 199 regions where we have confidence to combine the drug-evoked percent change readouts from the two imaging approaches. Second, considering only light sheet microscopy, regions were sampled at a denser rate, and we had more regions with at least 10 c-Fos+ cells in each brain across drug conditions. The number of regions that fulfill this criterion but was not in the combined set of 199 regions, was 97 regions. For these regions, we added to the data set by including only the light sheet microscopy data. Altogether, this yielded a data set of 296 regions for further analyses. The excluded regions were primarily tiny subcortical areas and sub-divisions of the cerebellum.

### Psilocybin and ketamine induce convergent and distinct differences in c-Fos expression across brain regions

**Figure 2** shows the percent difference in mean c-Fos positive cell density for either psilocybin or ketamine relative to saline for all 234 brain regions included in the analyses. The brain regions were sorted based on their membership in higher-order groupings (e.g., cortex, olfactory, hippocampus, etc.). This plot highlights the heterogenous effects of psilocybin and ketamine on c-Fos expression on a region-by-region basis. Broadly, regions in the cortex, thalamus, and brainstem systems had substantial differences, whereas regions in the olfactory and striatum / pallidum systems have relatively modest differences. The mean c-Fos cell count and mean drug-induced density change values are provided as spreadsheets in **Supplementary Table 2** and **Supplementary Table 3**. We also plotted the results using only serial two-photon imaging data (**Figure S1**) or only light sheet imaging data (**Figure S2**).

**Figure 2.**
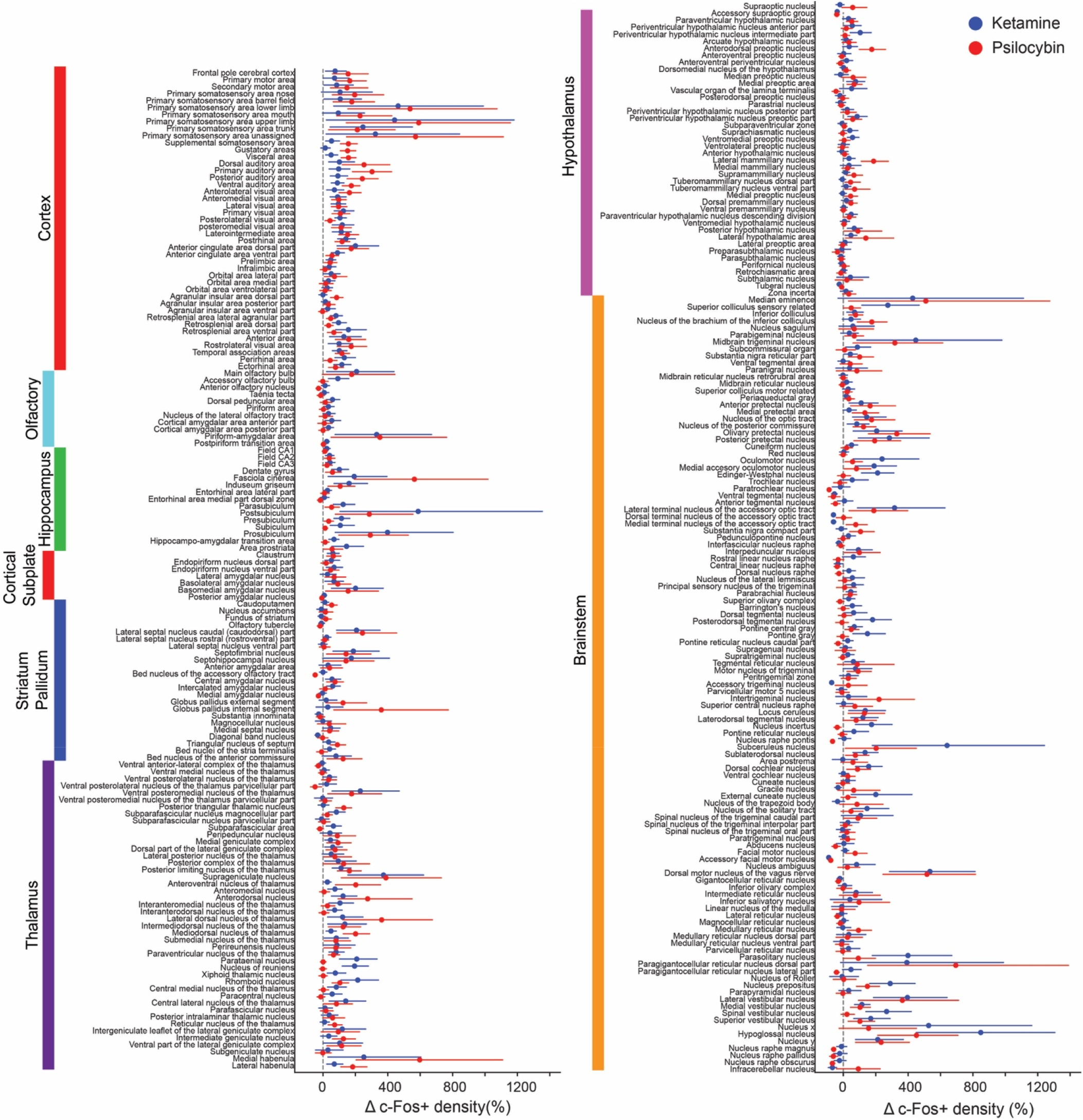
Effects of psilocybin and ketamine on regional c-Fos expression. Drug-evoked percent change in c-Fos+ cell density for psilocybin (red) and ketamine (blue). Circle, mean. Line, bootstrapped 95% confidence intervals assuming normal distribution.

To contrast effects of psilocybin and ketamine more clearly, we made a scatterplot of the average drug-induced c-Fos change by each drug (**Figure 3A** and **Figure S3**). For most brain regions examined, psilocybin and ketamine both increased c-Fos expression (upper right quadrant) or both decreased c-Fos expression (lower left quadrant, **Figure 3A**), although select locations showed preferential response to psilocybin or ketamine. We highlight several cortical regions of interest, either because of prior studies or because of large drug-evoked effects. In the medial frontal cortex, psilocybin and ketamine induced the largest change in c-Fos expression for the dorsal regions (i.e., dorsal anterior cingulate cortex (ACAd; **Figure 3B**) and ventral anterior cingulate cortex (ACAv)), with smaller increases as a function of depth for the more ventral regions (e.g., prelimbic area (PL). Psilocybin elicited greater elevation of c-Fos expression than ketamine in the dorsal agranular insular area (AId; **Figure 3B**). Conversely, ketamine evoked larger differences than psilocybin in the piriform (PIR) and field of CA1 (CA1); **Figure 3B**). Intriguingly, psilocybin and ketamine were both effective at elevating the immediate early gene levels across several areas in the visual hierarchy including: the primary visual area (VISp), the posterolateral visual area (VISpl), anterolateral visual area (VISal), posteromedial visual area (VISpm), and anteromedial visual area (VISam). Lastly, both ketamine and psilocybin increased c-Fos expression in retrosplenial cortical regions (RSPd, RSPv, RSPagl).

**Figure 3.**
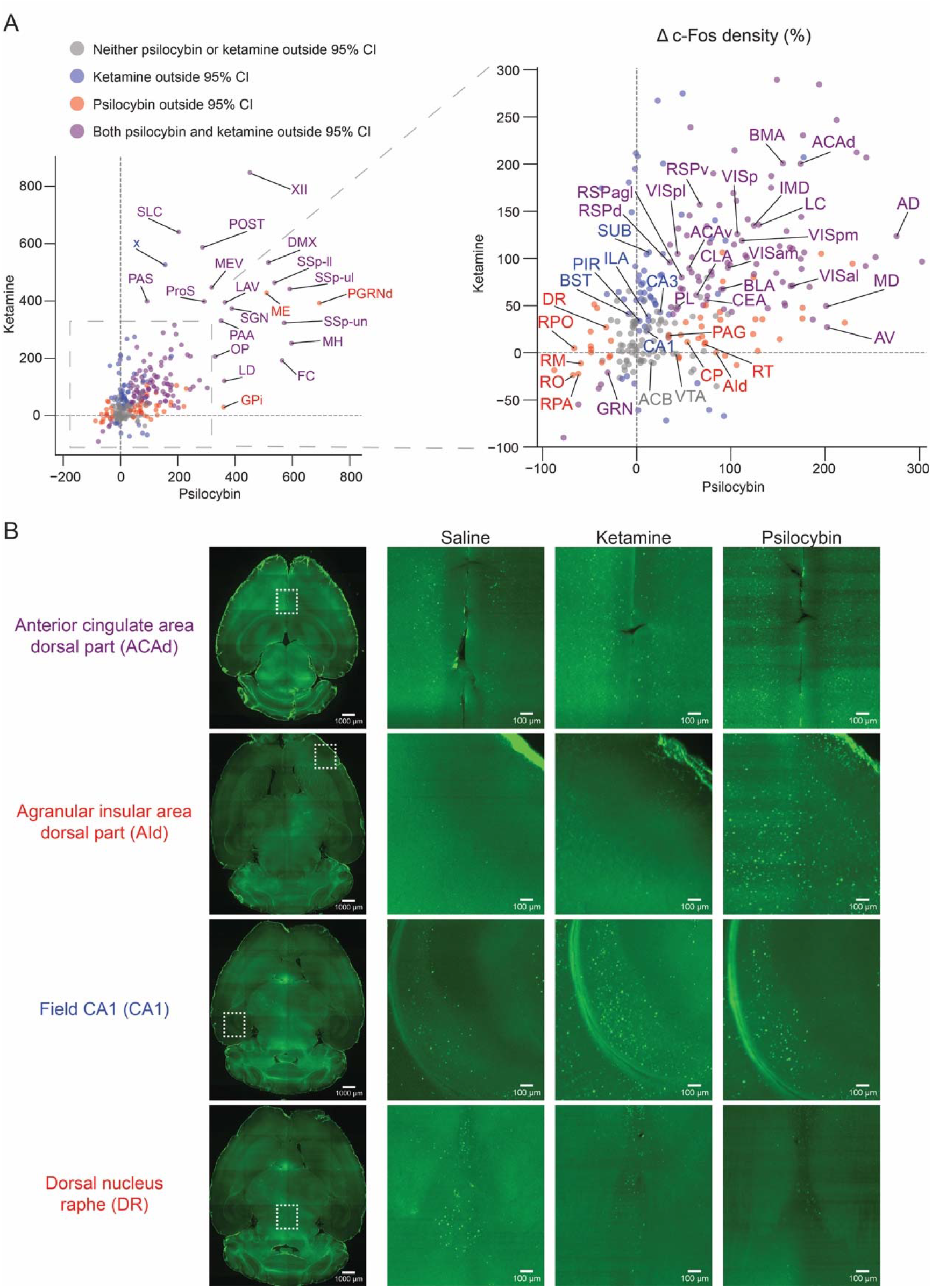
Common and distinct regions targeted for c-Fos expression by psilocybin and ketamine. **(A)** Scatter plot of mean drug-evoked percent change in c-Fos+ density for psilocybin (x-axis) versus ketamine (y-axis). For list of abbreviations, see Supplementary Table 3. **(B)** Example images from light sheet microscopy for select cortical and subcortical brain areas. Due to background intensity, for visualization purposes, we performed gamma correction on the magnified images, using the same adjustment for each row of images.

Psilocybin and ketamine also have shared and divergent targets in subcortical regions of interest. The locus coeruleus (LC) was notable for large raises in c-Fos expression following both psilocybin and ketamine. Similarly, the lateral and medial habenula had noted increases in c-Fos following both ketamine and psilocybin administration. The claustrum (CLA), several amygdalar (CEA, BLA, BMA) as well as the anterior (AV, AD) and midline thalamic nuclei (IMD, MD) also exhibited increased c-Fos expression following ketamine or psilocybin. The reticular nucleus of the thalamus (RT), the caudoputamen (CP), and periaqueductal gray (PAG) were more selectively targeted by psilocybin than ketamine. Conversely, the bed nucleus of the stria terminalis (BST) and key portions of the hippocampal circuit (SUB, CA1, CA3) were more selectively targeted by ketamine. However, not all regions exhibited increases in c-Fos positive cells: both drugs were effective at suppressing c-Fos expression in the Gigantocellular reticular nucleus (GRN) and psilocybin was selective for decreasing c-Fos expression in several raphe nuclei, in particular the dorsal nucleus raphe (DR; **Figure 3B**) as well as the nucleus raphe pontis (RPO), nucleus raphe magnus (RM), nucleus raphe pallidus (RPA), nucleus raphe obscurus (RO). Regions with non-significant change in c-Fos expression included the nucleus accumbens (ACB) and ventral tegmental area (VTA).

Several aspects of these results are consistent with prior work, which validate the whole-brain mapping approach. For instance, the medial frontal cortex, particularly the anterior cingulate cortex, is known to increase firing activity acutely following the systemic administration of N-methyl-D-aspartate receptor (NMDAR) antagonists ^17,53^, including subanesthetic ketamine ^54^. Our identification of increases of c-Fos signals in retrosplenial cortex is consistent with an earlier study using a slightly higher but still subanesthetic dose of ketamine ^55^ and a recent report of ketamine-evoked oscillatory activity in retrosplenial areas, especially in ventral regions ^56^. For psychedelics, a most telltale sign was the decrease in c-Fos expression in dorsal raphe and other raphe nuclei, which echoes the classic finding of near-complete cessation of spiking activity following various psychedelics including psilocin ^20,21,57^. However, many characterized regions in this study were previously underappreciated as potential mediators of psilocybin’s action. Effect for psilocybin in insular area, which is implicated in interoception and emotional awareness ^58^, and lateral habenula, which is maladaptively affected by stress and depressive state ^59–61^, are a few examples of potential interest.

### Receptors and cell types that may contribute to drug-induced c-Fos expression

To gain insight into the mechanisms by which these pharmacological agents act, we analyzed our c-Fos expression data in reference to publicly available atlas of gene expression. We leveraged the Allen Brain Institute’s *in situ* hybridization maps of the entire mouse brain ^62^, which has the mRNA transcript levels of all 19,413 genes in the mouse genome including various receptors and cell-type-specific markers. This allowed us to, for example, determine the relative expression levels of key serotonin receptor genes, including *Htr1a, Htr2a*, and *Htr2c*, in regions across the entire mouse brain (**Figure S4**). To estimate the relevance of each of the 19,413 genes, we correlated its regional expression levels with the regional drug-evoked c-Fos expression (**Figure 4A**). When the analysis was applied to the entire brain, psilocybin- and ketamine-induced differences in c-Fos did not correlate particularly well to many candidate receptors on a brain-wide scale (**Figure 4 B-E**), with the exception of several glutamate receptor genes for psilocybin-evoked expression including *Grin2a* and *Grin2b* (92^nd^ and 74^th^ percentiles of all genes), which encode the GluN2A and GluN2B subunit of NMDA receptors respectively (**Figure 4D**). Among serotonin receptor subtypes, psilocybin-induced differences in c-Fos expression was correlated best with *Htr2a* (63^th^ percentile), which encodes the 5-HT_2A_ receptor and consistent with the receptor being the primary driver of the acute hallucinogenic effects ^63^ and possibly the longer-term plasticity processes ^14,64^. The importance of the specific receptors was clearer when we restrict the analyses to only regions in the cortex (**Figure 4 F-I**). We found qualitatively similar results for psilocybin, with strong correlations for *Grin2a* (93^rd^ percentile), *Grin2b* (90^th^ percentile), and *Htr2a* (73^th^ percentile) (**Figure 4F, H**). The *Grin2a and Grin2b* genes also had a robust correlation (86^th^ and 83^rd^ percentiles) with ketamine-induced c-Fos expression in the cortex (**Figure 4I**), corroborating recent studies showing the importance of GluN2B for ketamine’s antidepressant action ^54,65^. Lastly, for the cortex, there are established genetic markers for various excitatory and inhibitory cell types ^66^. We found several genes that correlated well with both psilocybin- and ketamine-induced differences in c-Fos expression: *Pvalb* (99^th^ percentiles for both psilocybin and ketamine), a marker for GABAergic fast-spiking interneurons, *Lamp5* (96^th^ and 98^th^ percentiles), a marker for a subclass of GABAergic interneurons including neurogliaform cells and single bouquet cells, and *Fezf2* (74^th^ for both), a marker of extra-telencephalic projecting layer-5 pyramidal cells that include pyramidal tract neurons (**Figure 4F, G**). Cumulatively, this exploratory analysis suggests that NMDA receptor distribution is predictive of both psilocybin- and ketamine-evoked c-Fos expression patterns, particularly in the cortex, and therefore is a clue to support glutamatergic signaling as a potential convergent mechanism that shape the effects of psilocybin and ketamine on neural plasticity.

**Figure 4.**
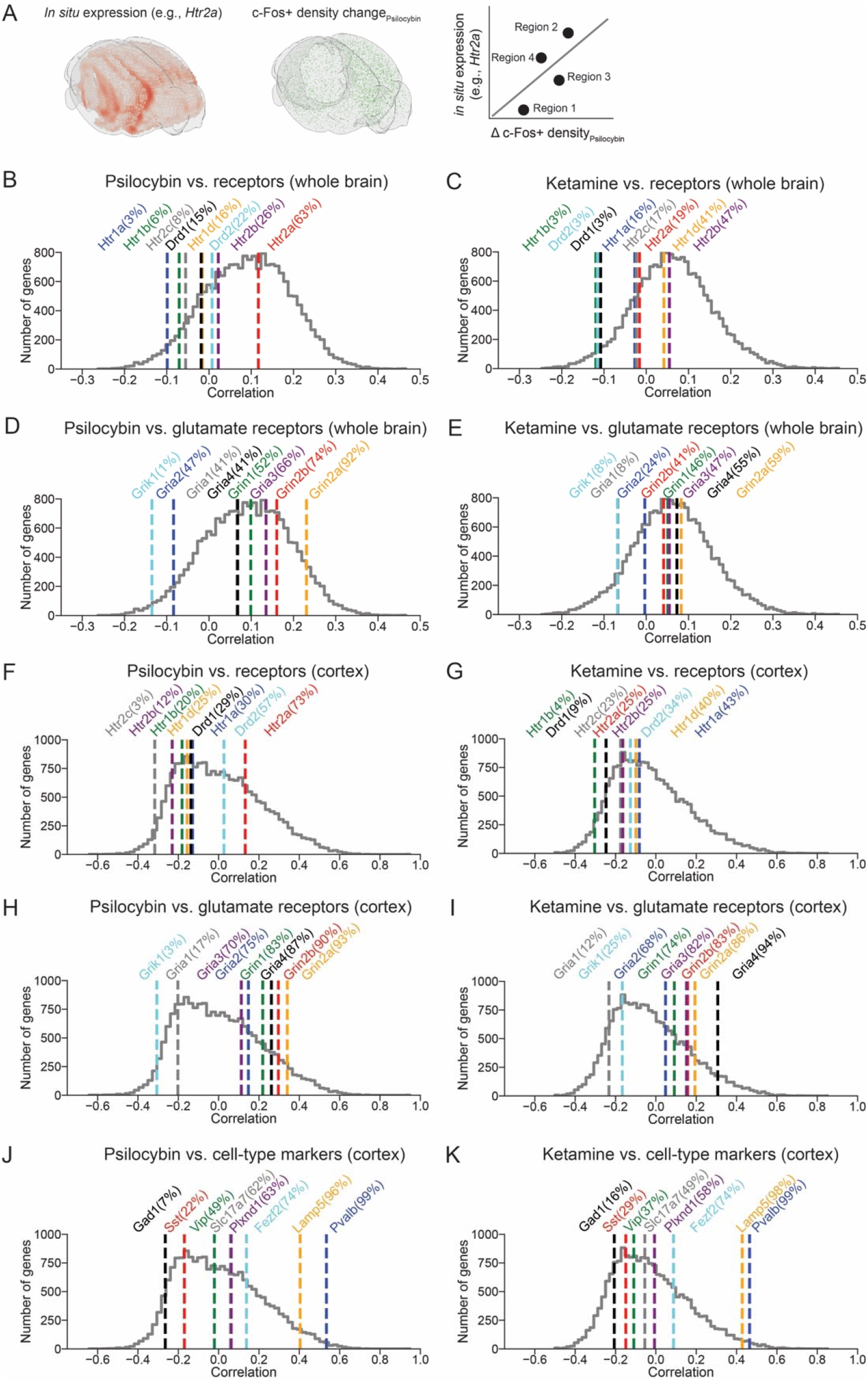
Potential receptors and cell types contributing to drug-evoked c-Fos expression. **(A)** Schematic illustrating the analysis procedure. The mRNA transcript levels of a particular gene (e.g., *Htr2a*) (left, interpolated from sagittal sections to yield 3D rendering using Brainrender ^103^) was compared with drug-evoked percent change in c-Fos+ density (middle), on a region-by-region basis to calculate a correlation coefficient (right). **(B)** Correlation coefficients computed for psilocybin condition using regions across the entire brain. Colored lines, correlation coefficients for select serotonin and dopamine receptor genes, with percentile indicated within the parenthesis. Grey line, histogram of correlation coefficients for all 19,413 genes in the mouse genome. **(C)** Similar to (B) for ketamine. **(D)** Similar to (B) for glutamate receptors. **(E)** Similar to (D) for ketamine. **(F - I)** Similar to (B - E) except using only regions within the cortex. **(J, K)** Similar to (F, G) for major cell-type marker genes.

## DISCUSSION

Our study revealed the similarities and differences in the expression of the immediate early gene c-Fos following administration of psilocybin and ketamine. The systematic, unbiased mapping approach provides a comprehensive coverage of all brain regions and should be a valuable resource for the community seeking to understand the effects of these compounds. For psilocybin, the data not only affirm the likely importance of well-studied brain regions such as the anterior cingulate cortex and dorsal raphe, but also pinpoint several underappreciated regions such as reticular nucleus of the thalamus and insular cortex that may be crucial for drug action. Furthermore, correlation of drug effects with cell-type markers and receptor genes highlighted glutamatergic receptors not only for ketamine, but intriguingly also for psilocybin. The results suggest that glutamatergic receptors may be a potential convergent target for how psilocybin and ketamine produce their long-lasting effects on neural plasticity.

We chose c-Fos for this study because it is a well-characterized immediate early gene and, unlike other plasticity-related genes such as Arc, the nuclear staining of c-Fos is amenable to automated cell counting using machine learning tools. However, there are several limitations. First, the use of transgenic animals may yield signals that differ from the endogenous c-Fos expression. Such discrepancy was documented in a recent comparison following whisker learning in mice ^50^, and may due in part to numerous enhancers surrounding the c-Fos gene being important for response to stimuli ^49^. This caveat is alleviated in part in this study by also studying c-Fos expression in wild type mice using light sheet microscopy. Second, c-Fos captures activity-dependent transcription in the nucleus, but drug-evoked neural plasticity is likely to also rely on local mechanisms, such as local protein synthesis in the dendritic compartments. Third, although c-Fos expression increases are widely thought to reflect elevated spiking activity, the relationship remains unclear. Our results broadly support this view, with ketamine’s effect on anterior cingulate cortex and psilocybin’s effect on dorsal raphe consistent to prior electrophysiological measurements. However, there are also discrepancies: for example, we observed psilocybin-induced c-Fos expression in medial frontal cortex and primary visual cortex, but studies indicate that the overall effects of psychedelics, at least for the phenethylamine DOI, on spiking activity should be suppressive in these regions ^17,67^. Finally, as an immediate early gene, the expression of c-Fos is expected to evolve over time. We have chosen a single time point to capture peak expression level, but future studies may include additional measurements to delineate a greater time course ^68^.

Ketamine is primarily a NMDAR antagonist, and therefore has direct action on glutamatergic receptors in the brain. There is growing consensus that the therapeutic effects of ketamine depend critically on its influence on glutamatergic signaling ^69–72^. By contrast, psychedelics such as psilocybin are serotonin receptor agonists, and therefore studies of the drugs’ effect on neural plasticity have mostly focused on the pharmacology of serotonin ^12–14,64^. The exploratory analyses in this study revealed a correlation between *Htr2a* transcript levels and psilocybin-evoked c-Fos expression, consistent with the known receptor interaction. But we also observed robust relationships for glutamatergic receptors including *Grin2a* and *Grin2b,* particularly for cortical regions, which means that the presence of glutamatergic receptors is an even stronger indicator for whether a cortical region is sensitive to psilocybin-evoked neural plasticity. The results therefore provide empirical support for an interplay between serotoninergic and glutamatergic signaling for psilocybin’s plasticity effects, which has been speculated before as a potential convergent mechanism between ketamine and psychedelics ^9,10.^

Looking forward, the approach used here could be extended to study other drugs and new chemical entities. This may be other psychedelics, which include a large array of compounds ^1,2^ that vary in their binding affinity to various serotonin and non-serotonin receptors ^73^. The effects of psilocybin and ketamine may be compared to other antidepressant agents, such as brexanolone ^74^ and lumateperone ^75^, and new treatment options, such as other glutamate-targeting drugs ^76,77^ or nitrous oxide ^78^. Understanding the shared and disparate mechanisms underlying contrasting drugs will be crucial in developing a greater understanding of the pharmacology of rapid-acting antidepressants.

## ACKNOWLEDGMENTS

We thank Neil Savalia, Adam Tyson, and Boris Heifets for discussions about the analysis. Psilocybin was generously provided by Usona Institute’s Investigational Drug & Material Supply Program; the Usona Institute IDMSP is supported by Alexander Sherwood, Robert Kargbo, and Kristi Kaylo in Madison, WI. This work was supported by the Yale Program in Psychedelic Science, NIH/NIMH grant R01MH121848 (A.C.K.), NIH/NIMH grant R01MH128217 (A.C.K.), One Mind – COMPASS Rising Star Award (A.C.K.), and NIH/NIGMS medical scientist training grant T32GM007205 (P.A.D.).

## AUTHOR CONTRIBUTIONS

P.A.D. and A.C.K. designed the experiments. P.A.D. and L.-X. S. prepared the samples. P.A.D. conducted the experiments and analyzed the data. P.A.D. and A.C.K. wrote the paper.

## METHODS AND MATERIALS

### Animals

Equal numbers of male and female animals were used for the study. Animals were randomly assigned to the saline, ketamine, or psilocybin condition. No animals were excluded from data analysis. Adult, 6 to 20-week-old *cfos^GFP^* mice ^45,46^ (B6.Cg-Tg(Fos-tTA,Fos-EGFP*)1Mmay/J, #018306, The Jackson Laboratory) were used for the serial two-photon whole-brain mapping experiments. Adult, 8-week-old C57BL/6J mice (#00064, The Jackson Laboratory) were used for the light sheet whole-brain mapping experiments. All animals were housed and handled according to protocols approved by the Institutional Animal Care and Use Committee (IACUC) at Yale University.

### Serial two-photon microscopy – sample preparation and imaging

The *cfos^GFP^* mice were injected with either saline (10 mL/kg, i.p.), ketamine (10 mg/kg, i.p.), or psilocybin (1 mg/kg, i.p.). At 3.5 hours after the injection, the mice were deeply anesthetized with isoflurane and transcardially perfused with phosphate buffered saline (P4417, Sigma-Aldrich) followed by paraformaldehyde (PFA, 4% in PBS). The brains were fixed in 4% PFA for 12 hours at 4°C. Brains were transferred to PBS with 0.1% sodium azide until they were sectioned and imaged. Whole-brain serial two-photon tomography imaging was performed using the previously described TissueCyte 1000 system ^79^. Briefly, brain samples were imaged using a laser with an excitation wavelength of 920 nm, and emitted fluorescence was captured across three channels (channel 1: 560–680, channel 2: 500–560, and channel 3: 400–500 nm). GFP fluorescence was detected in channel 2. Autofluorescence signals were detected in channel 1 and 3. Approximately 140 serial block-face images were acquired at 100-μm spacing for each brain at 1.4 μm/pixel XY sampling. The imaging steps were done blinded to the treatment conditions at TissueVision, Inc (Newton, MA).

### Serial two-photon microscopy – analysis

Tiled brain images were processed through the QUINT workflow ^80^ for registration and quantification of GFP-expressing, c-Fos-positive (c-Fos+) cells. First, images were registered to the Allen Brain Atlas (Allen Reference Atlas – Mouse Brain [Adult Mouse]. Available from https://atlas.brain-map.org) using the autofluorescence signals and QuickNII tool ^81^ was used to guide a rigid, affine registration and map brain slices into three-dimensional space based on key anatomical landmarks. Next, the VisuAlign tool (RRID:SCR_017978) was used to further improve the registration using nonlinear refinements. The c-Fos+ cells in each image were segmented with two levels of classification. An initial pixel-level classification and then an object-level classification performed via supervised machine-learning with *ilastik*^82^. Lastly, the registered tiled brain images were overlaid with the segmented c-Fos+ cells using the Nutil tool ^83^ to determine the count of c-Fos+ cells in each region.

### Light sheet microscopy – sample preparation and imaging

C57BL/6J mice were injected with either saline (10 mL/kg, i.p.), ketamine (10 mg/kg, i.p.), or psilocybin (1 mg/kg, i.p.). At 2 hours after the injection, the mouse was deeply anesthetized with isoflurane and transcardially perfused with phosphate buffered saline (P4417, Sigma-Aldrich) followed by paraformaldehyde (PFA, 4% in PBS). The brains were fixed in 4% PFA for 24 hours at 4°C. Brains were then transferred to PBS with 0.1% sodium azide until brain clearing and labeling. Whole mouse brains were processed following the SHIELD protocol ^84^. Samples were cleared for 4 days at 42°C with SmartClear (LifeCanvas Technologies), a device employing stochastic electrotransport ^85^. Cleared samples were then actively immunolabeled using eFLASH technology integrating stochastic electrotransport ^85^ and SWITCH ^86^. Each brain sample was stained with primary antibodies, 3.5 μg of rabbit anti-c-Fos monoclonal antibody (Abcam, #ab214672), and 10 μg of mouse anti-NeuN monoclonal antibody (Encor Biotechnology, #MCA-1B7) followed by fluorescently conjugated secondaries in 1:2 primary:secondary molar ratios (Jackson ImmunoResearch). After active labeling, samples were incubated in EasyIndex (LifeCanvas Technologies) for refractive index matching (n = 1.52) and imaged at a magnification of 3.6X with a SmartSPIM light sheet microscope (LifeCanvas Technologies) at 1.8 μm/pixel XY sampling with 4 μm Z sampling over the entire brain. The imaging steps were done blinded to the treatment conditions at LifeCanvas Technologies (Cambridge, MA).

### Light sheet microscopy – analysis

Images were tile corrected, de-striped, and registered to the Allen Brain Atlas using an automated process. Specifically, a NeuN channel for each brain was registered to 8-20 atlas-aligned reference samples, using successive rigid, affine, and b-spline warping algorithms with SimpleElastix ^87^. An average alignment to the atlas was generated across all intermediate reference sample alignments to serve as the final atlas alignment value for the individual sample. Automated cell detection was performed using a custom convolutional neural network through the TensorFlow python package ^88^. The cell detection was performed by two networks in sequence. First, a fully-convolutional detection network ^89^ based on a U-Net architecture ^90^ as used to find possible locations of c-Fos positive cells. Second, a convolutional network using a ResNet architecture ^91^ was used to classify each location as positive or negative hit. Using the atlas registration, each cell location was projected onto the Allen Brain Atlas to quantify the number of fluorescent c-Fos+ cells for each atlas-defined region.

### Bridging serial two-photon and light sheet imaging data

The Allen Mouse Brain Common Coordinate Framework (Allen CCF) contains over one thousand brain region delineations that are arranged hierarchically ^51^. To constrain our results, we focus our analysis here on the 316 ‘summary structures’ as proposed by the Allen CCF authors ^51^. To bridge the data from serial two-photon imaging and light sheet imaging, we had a two-step procedure. First, whenever possible, we used data from both imaging methods. Some small brain regions had very few cells, which could inflate drug-evoked changes in c-Fos+ cell density. We identified 234 brain regions where ≥10 c-Fos+ cells were detected in each of the brains across all drug treatment conditions by both imaging methods. Then we determined the drug-evoked change in c-Fos+ density change for psilocybin. For 35 regions, the two imaging methods yielded divergent results exceeding a six-fold difference, presumably due to the differences in axial sampling step size for serial two-photon imaging (100 μm) versus light sheet microscopy (4 μm). This yields 199 brain regions in which we are confident the two imaging methods produce comparable results. We combine the data by considering the drug-evoked change in c-Fos+ density in samples obtained by both methods. Second, light sheet microscopy has denser sampling. There were 97 regions in which ≥10 c-Fos+ cells were detected in each of the brains across all drug treatment conditions from light sheet microscopy but were not part of the 199 brain regions in the combined data set. We added these to the data set by including only the drug-evoked change in c-Fos+ density in samples obtained by light sheet microscopy. The total data set thus contains 296 regions.

### *In situ* hybridization

We accessed publicly available *in situ* hybridization data of all mouse genes across the entire brain ^62^ to assess the relative expression of each gene in each brain region via custom code through the AllenSDK ^92,93^. We used the regional density of RNA expression to quantify the expression of every gene in each brain region of interest. For each gene, we further calculated the Pearson correlation coefficient between its regional expression levels with regional drug-induced differences in c-Fos expression.

## Data and code availability

The data that support the findings and the code used to analyze the data in this study will be made publicly available at https://github.com/Kwan-Lab. The data for relative expression of all gene in each brain region will be shared at https://alexkwanlab.org. Raw image files are terabytes in size and will be available upon request.

The following open-source software was used: Python, conda, Numpy ^94^, SciPy ^95^, IPython ^96^, seaborn ^97^, Matplotlib ^98^, Pandas ^99^, xarray ^100^, statsmodels ^101^, allenCCF ^102^, brainrender ^103^, and Jupyter notebook ^104^. We are grateful to the creators and maintainers of these open-source tools.

## SUPPLEMENTARY MATERIALS

**Supplementary Table 1:** The brain region in the analysis. “Region_Name”, “Region_ID”, “abbreviation” and “Brain Area” follow the conventions set by Allen Institute for Brain Science. “Volume” is the mean volume determined for each region using serial two-photon microscopy in this study. “Region_index” is the index used to plot Figure 1J.

**Supplementary Table 2:** Mean density of c-Fos+ cells (cells/mm^3^) per brain region for saline, ketamine, and psilocybin.

**Supplementary Table 3:** Mean percent change and 95% confidence intervals of density of c-Fos+ cells per brain region for psilocybin and ketamine, relative to saline.

**Figure S1:**
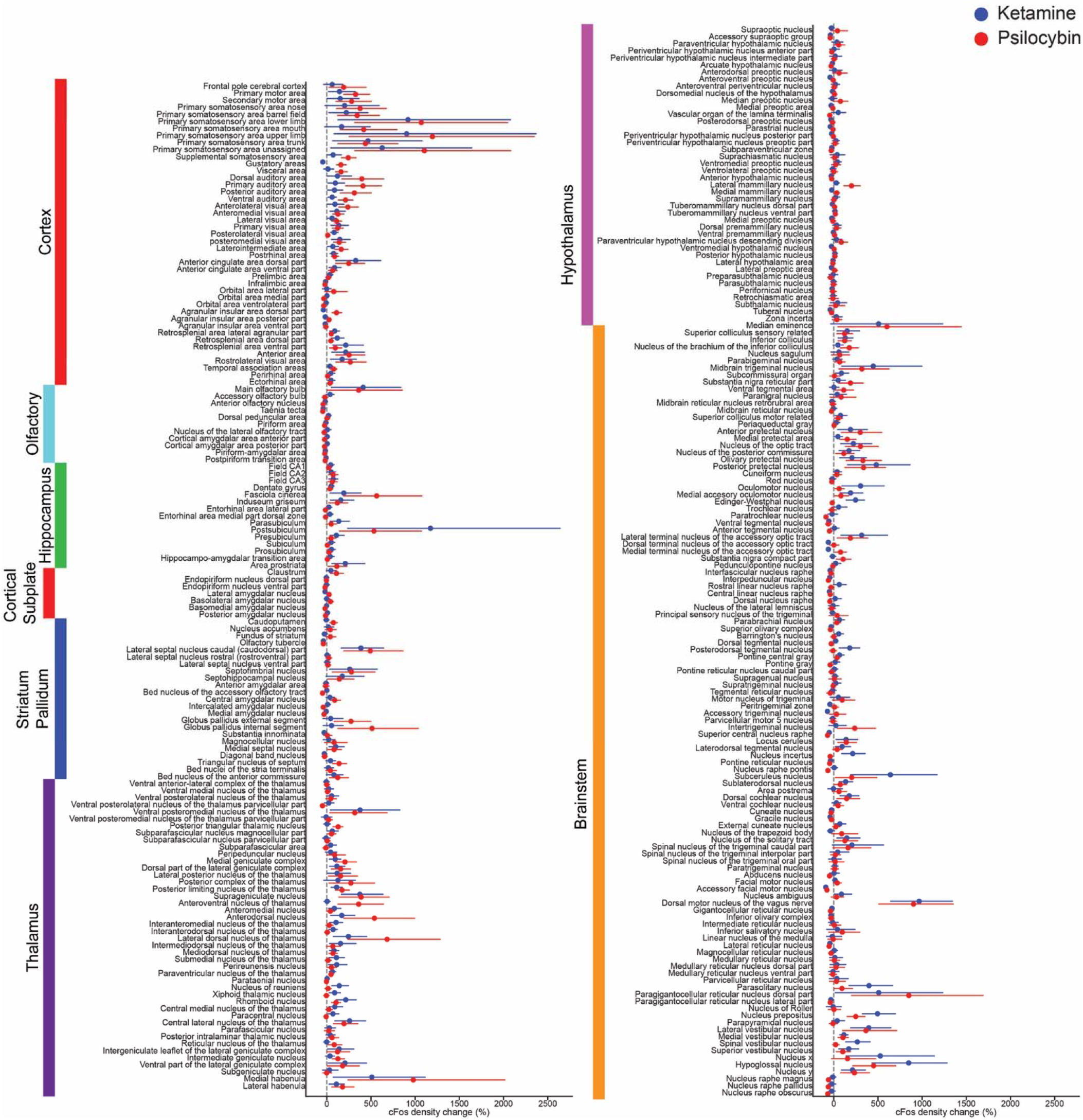
Effects of psilocybin and ketamine on regional c-Fos expression as measured by serial two-photon microscopy. Percent change in c-Fos density from saline vehicle baseline for psilocybin (red) and ketamine (blue). Circle, mean. Line, bootstrapped 95% confidence intervals assuming normal distribution.

**Figure S2:**
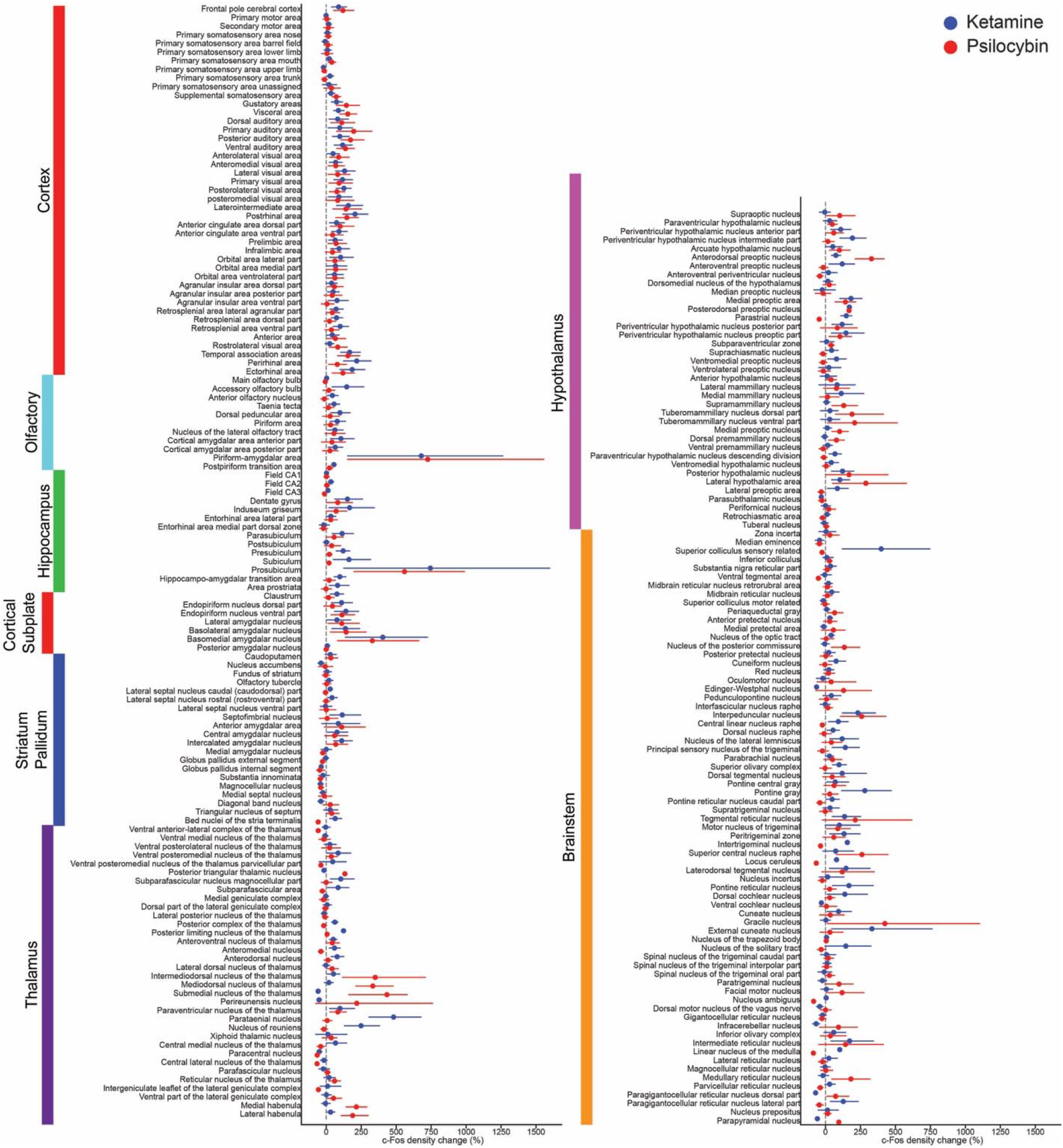
Effects of psilocybin and ketamine on regional c-Fos expression as measured by light sheet microscopy. Percent change in c-Fos density from saline vehicle baseline for psilocybin (red) and ketamine (blue). Circle, mean. Line, bootstrapped 95%confidence intervals assuming normal distribution.

**Figure S3:**
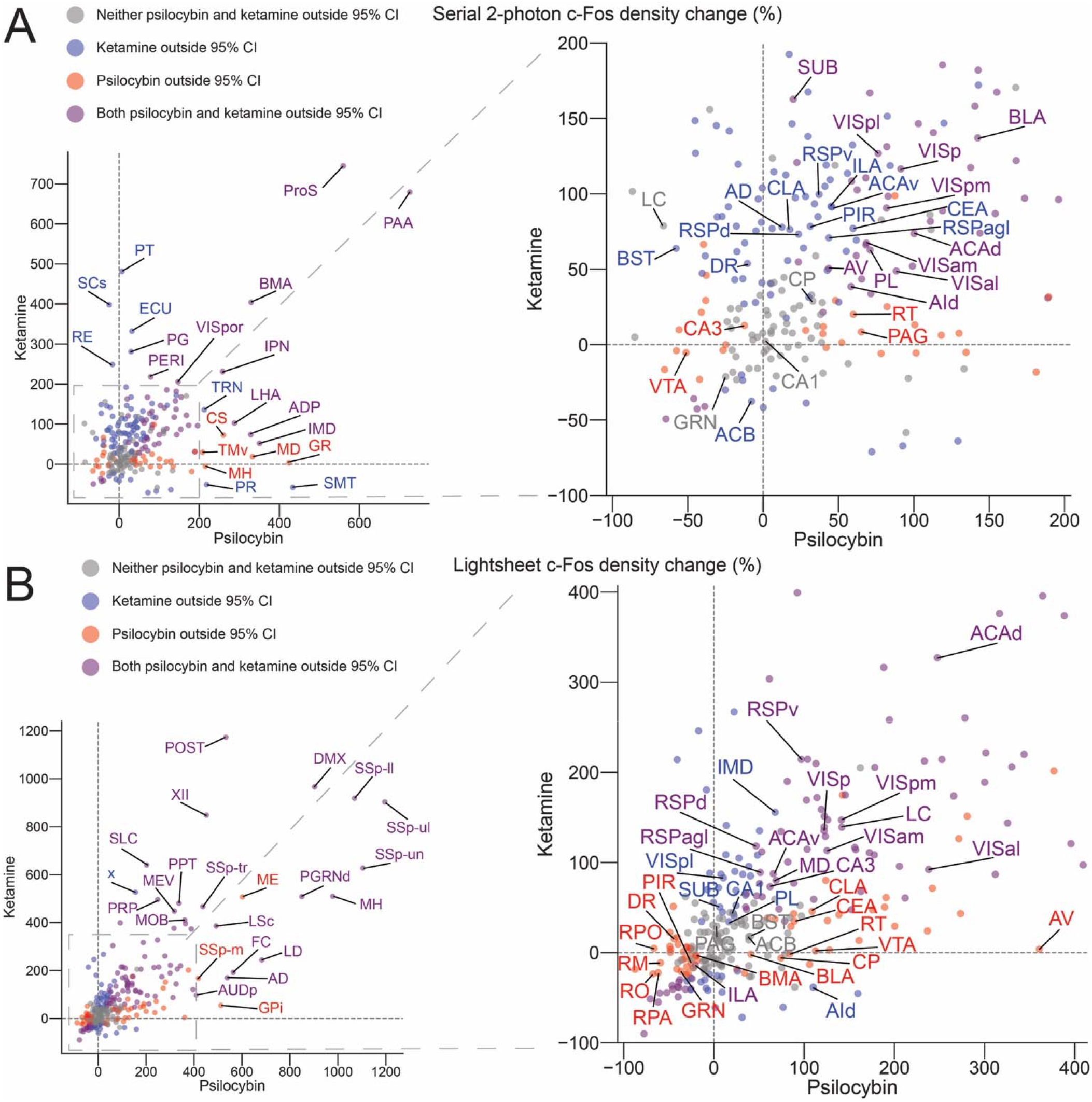
Psilocybin and ketamine induce convergent and distinct differences in c-Fos expression. **(A)** Scatter plot of mean percent change in c-Fos density from saline vehicle, for psilocybin (x-axis) versus ketamine (y-axis) as measured by serial two-photon microscopy. **(B)** Similar to (A) but as measured by light sheet microscopy.

**Figure S4:**
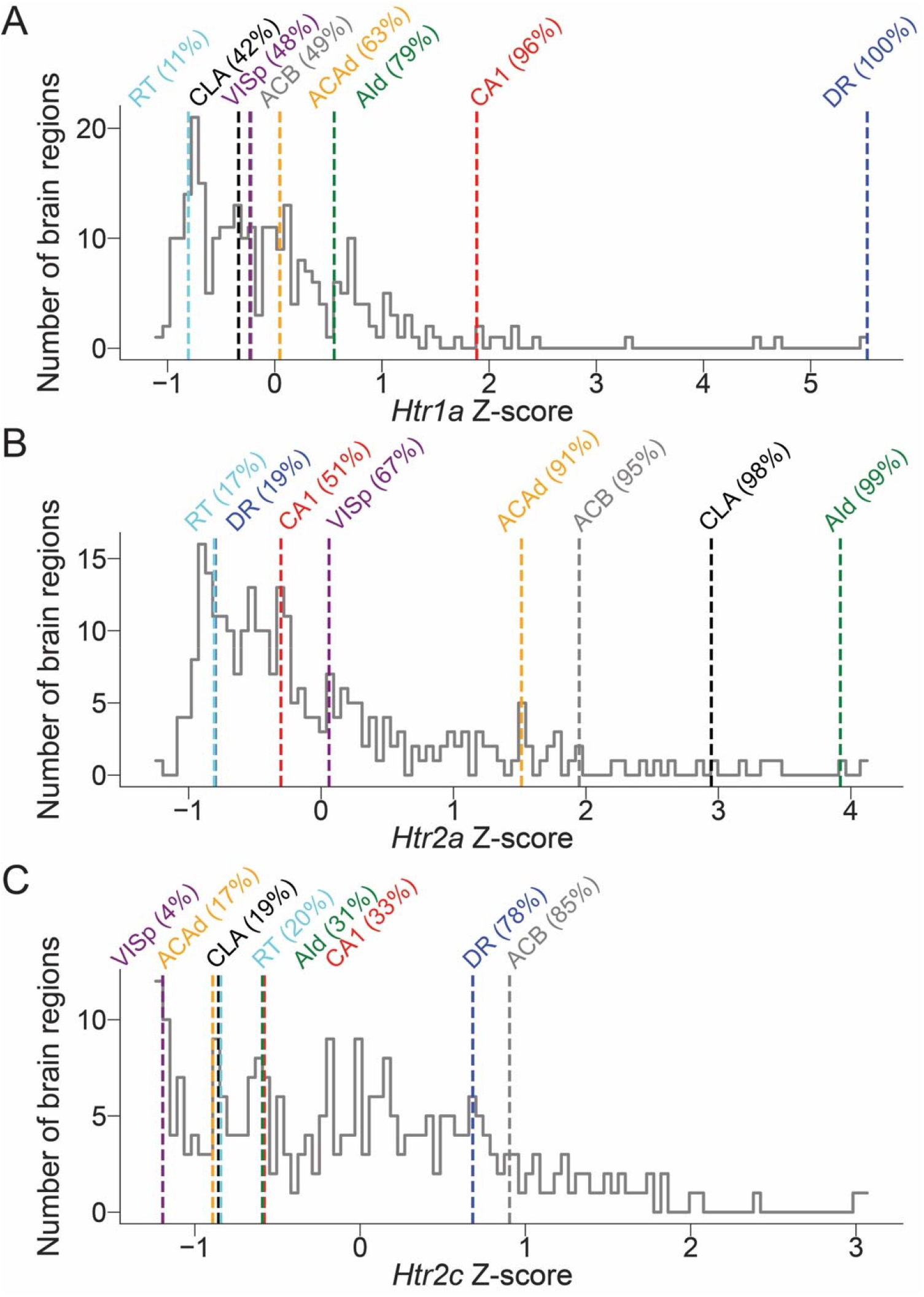
Differential expression of serotonin receptors across brain regions. **(A)** Relative gene expression levels of *Htr1a* gene from Allen Institute *in situ* hybridization database across all brain regions. Grey line, histogram of Z-score for *Htr1a* gene for all brain regions. Colored lines, Z-score and percentile rank for specific brain regions. **(B)** similar to **(A)** but for *Htr2a* gene. **(C)** similar to **(A)** but for *Htr2c* gene.

